# The phoretic mite (Arachnida: Acari) assemblage of the Douglas-fir beetle, *Dendroctonus pseudotsugae* Hopkins (Coleoptera: Curculionidae)

**DOI:** 10.1101/2023.06.16.545383

**Authors:** Laura-Anne Browning, Dezene P.W. Huber

## Abstract

The phoretic mite assemblage of the Douglas-fir beetle, *Dendroctonus pseudotsugae* Hopkins (Coleoptera: Curculionidae) has not been as thoroughly documented as that of some other ecologically and economically important bark beetle species. Phoretic mites can impact individual fitness and population dynamics of their hosts and documenting the mite assemblage associated with a bark beetle may provide information on their ecological and interactive roles. We caught Douglas-fir beetles over two summers in central British Columbia, Canada and sorted the associated mites into morphospecies. Representatives of the morphospecies were DNA barcoded (cytochrome oxidate I barcode region) which indicated at least nine distinct Operational Taxonomic Units (OTUs). There was a mean of 50.5 ± 4.7 mites per beetle with both females and males carrying similar numbers of most mite species. However one OTU (Sarcoptiformes: Hemisarcoptidae) was found in substantially higher numbers than all other OTUs, and was always clustered in large aggregations in an anterior pocket on the sub-elytral surface. When B1 was removed from the mean, there were only 1.3 ± 0.2 mites per beetle. The consistent high numbers of that OTU in conjunction with its consistent anatomical aggregation suggests an important interaction between that mite species and the Douglas-fir beetle.

## Introduction

Phoresy is a dispersal strategy where the dispersing organism (the phoront) associates with another organism (the host) to move between habitats. Many mite groups, which are small, wingless, and often associated with ephemeral resources, are dependent on a phoretic lifestyle with larger, more mobile organisms. The development of life stages with anatomical and other modifications specialized for phoresy is common in mite taxa (Bartlow and Agosta 2021; Walter and Proctor 2013). Phoretic mites form complex interactions with their hosts as well as with the other symbionts associated with their hosts and can ultimately impact population dynamics of their hosts.

For example, some mite species that form phoretic relationships with the southern pine beetle, *Dendroctonus frontalis* Zimmerman (Coleoptera: Curculionidae) are also predators of the beetle larvae and pupae and have been identified as potential biological control agents (Moser 1975). Predatory mites may also benefit their phoretic hosts by feeding on the natural enemies of these hosts (Hofstetter et al. 2023). Other *Tarsonemus* spp. G. Canestrini & Fanzago (<u>Trombidiformes</u>: <u>Tarsonemidae</u>) mites have structures called sporothecae that carry ascospores of fungi that compete with fungi carried in its bark beetle host’s mycangia that are beneficial to beetle’s larvae – the presence of this mite may result in population declines in southern pine beetles (Hofstetter et al. 2006; Lombardero et al. 2003; Moser 1985). Other studies suggest that mite loads influence the flight and mobility of their beetle hosts (Atkins 1960; Rocha et al. 2009).

The mite assemblage of the Douglas-fir beetle, *Dendroctonus pseudotsugae* Hopkins (Coleoptera: Curculionidae), remains poorly documented as most studies on bark beetle phoronts focus on more economically important species like *D. frontalis* and *Ips typographus* L. (Coleoptera: Curculionidae) (Hofstetter et al. 2014; Hofstetter et al. 2015). The Douglas-fir beetle mainly attacks Douglas-fir trees, *Pseudotsuga menziesii* (Mirb.) Franco (Pinaceae), and occasionally western larch, *Larix occidentalis* Nutt. (Pinaceae) (Kelley and Farrell 1998). As such, the Douglas-fir beetle’s distribution coincides with the range of its host trees (Ruiz et al. 2010). Adult Douglas-fir beetles create galleries and lay eggs in the phloem of their host trees where offspring feed and develop into adults (Stark 1982). Douglas-fir beetles also inoculate the trees they attack with fungi that, among other things, block defenses of the host tree and can lead to tree death at high enough beetle abundances (Raffa 2008). The Douglas-fir beetle is a major economic concern across its range as it is sometimes capable of killing large numbers of hosts during outbreaks. For instance, in 2021 the Douglas-fir beetle was the third-most active bark beetle in British Columbia, affecting over 106,000 ha of forested land in the province (Ministry of Forests 2021). Irruptive population growth and epidemic-level outbreaks can also occur in combination with other forest disturbances and Douglas-fir beetle infestations may become more extensive with climate change (Bentz et al. 2010; Cole et al. 2022). By documenting the phoretic mite assemblage of the Douglas-fir beetle, we will gain a greater understanding of the biodiversity of organisms associated with this species and provide a basis for further study on their role in the population dynamics of the insect.

In this study, we used a combination of morphospecies and genetic analyses to describe the taxonomic diversity of the phoretic mite assemblage of the Douglas-fir beetle. We also examined mite loads in relation to host beetle sexes and emergence phenology. To our knowledge, this study is one of the first performed on the phoretic mite assemblage of Douglas-fir beetles.

## Methods

We sampled Douglas-fir beetles from May to August 2021 and 2022 in forested areas around the University of Northern British Columbia (UNBC, Prince George, British Columbia, 53.8922° N, 122.8134° W). The UNBC campus is situated in the sub-boreal spruce (SBS) zone and the dry warm subzone (SBSdw2) of British Columbia, which is characterized by mixed stands of lodgepole pine [*Pinus contorta* Douglas ex Loudon (Pinaceae)], Douglas-fir, and interior hybrid spruce [*Picea engelmannii* f. *glauca* (R.Sm.) Beissn (Pinaceae)]. In both years, 12-unit Lindgren multiple funnel traps (Lindgren 1983) baited with a commercially available Douglas-fir beetle pheromone trap blend (Synergy Semiochemicals Corporation, Burnaby, British Columbia) were used for sampling. In 2021, two traps were placed at each of three sites and no pesticide was used in the trap cups. In 2022, pest strips were required to stop the predation of Douglas-fir beetles by Cleridae and to gather an accurate count of the beetles captured in each trap. However, we observed that the contact-insecticide cubes (cut from Vapona No-pest Strip, Fisons Horticulture Inc. Mississauga, Ontario, Canada) killed some Douglas-fir beetles with their elytra spread open, which could have resulted in the loss of mites attached to the subelytral surface. Therefore, in 2022, two traps were placed at seven locations around UNBC – one trap at each site contained a small pesticide-infused polymer block while the other did not. Beetles collected from traps containing pesticide were counted to determine emergence phenology, while the beetles collected from traps not containing pesticide were inspected for mites. In both years, the traps were checked frequently (two to four times per week), and all live beetles from which mites were sampled were individually collected in microcentrifuge tubes containing ∼1.5 ml 95% ethanol to decrease the likelihood of including mites associated with other insects in our collection. Dead beetles in the traps containing pesticide in 2022 were collected into sealable plastic bags.

The sex of the beetles was determined through the presence or absence of the dorsal abdominal stridulating organ males use to create acoustic signals (Lyon 1958). Following the confirmation of the sex, the elytra were removed for inspection, and the entire body of each beetle was inspected for mites, which were then removed, counted, and categorized into morphospecies by features such as size, body shape, unique features, and colouration. Mites suspended in the ethanol were included in the count as beetles that were inspected for mites were collected individually into ethanol, so mites in the ethanol were likely associated with the beetles. Each mite was stored separately from its corresponding beetle host (also stored individually) in 95% ethanol. Representative mites of each morphospecies were chosen for DNA barcoding.

A total of 94 mites associated with Douglas-fir beetles were successfully sequenced (cytochrome oxidase 1 DNA barcoding region) at the Biodiversity Institute of Ontario (BIO, University of Guelph) and sequence data were used to group them into Operational Taxonomic Units (OTUs). Sequences >400 bp and with no contaminants or stop codons were considered successful and used in the analysis. A total of 33 beetle specimens were also barcoded at BIO, and all of them matched to existing Douglas-fir beetle sequences. All mite and beetle specimens were vouchered at BIO and barcode data are accessible in dataset DS-#####. OTU clustering was performed using the Barcode of Life Database (BOLD) dashboard with the BOLD aligner using MUSCLE (Edgar 2004, Ratnasingham and Hebert 2007). A maximum likelihood tree for the mites was constructed with MEGA using one sequence per OTU cluster to visualize a representative tree (Stecher et al. 2020).

The prevalence (proportion of beetles carrying each OTU) and abundance (mean number of a specific OTUs per beetle) of each mite OTU were recorded. A Spearman’s rank correlation test was used to assess the association between the number of beetles caught in the traps and the number of mites. This analysis was only performed for the samples caught until 10 July 2022 as the majority of beetle flight was captured in that time period. OTUs A-C were used in this analysis as only they were abundant enough to be found across most of the trapping season. A Mann-Whitney U test was used to assess if the number of associated OTUs A-F and H carried by each sex differed significantly between male and female host beetles. Non-parametric tests were used in this study as the data did not follow a bivariate normal distribution All statistical analyses were performed in Rstudio version 2022.12.0+353 with an alpha level of 0.05.

## Results

We counted a total of 10,047 mites from 137 beetles in 2021 and 7,045 mites from 141 beetles in 2022. We identified ten morphospecies that were found attached either underneath the elytra or on the ventral surface of the thorax (morphospecies A-J, Table 1). In three instances (morphospecies H – J), the mites were only found in the storage ethanol, but were included in the analysis for completeness. DNA barcoding distinguished nine OTUs (>2% sequence divergence), differing slightly from the morphospecies analysis but mostly supporting the morphological groupings. DNA barcoding repeatedly failed for morphospecies G, I and J, while morphospecies B was found to include two distinct OTUs (hereafter referred to as OTU B_1_ and B_2_) and their counts were separated in 2022. Morphospecies D was found to be two distinct, but related OTUs (hereafter referred to as OTU D_1_ and D_2_) but their counts were not separated in 2022 due to substantial morphological similarity that made distinguishing between them unreliable. Only counts from OTUs confirmed by DNA barcoding have been included in the prevalence, abundance, and phenological data analyses.

**Table 1.** Mite morphospecies associated with the Douglas-fir beetles and the locations that they were found on the insect hosts.

| Morphospecies | Attachment location |
| --- | --- |
| A | Under elytra |
| B<sub>1</sub> | Under elytra |
| B<sub>2</sub> | Under elytra |
| C | Under elytra |
| D<sub>1</sub> | Ventral thorax |
| D<sub>2</sub> | Ventral thorax |
| E | Under elytra |
| F | Ventral thorax |
| G | Under elytra |
| H | N/A (in ethanol) |
| I | N/A (in ethanol) |
| J | N/A (in ethanol) |

The barcoded specimens belonged to three Orders: Mesostigmata (OTUs A, C, D_1_, D_2_, and E), Trombidiformes (OTUs B_2_ and F) and Sarcoptiformes (OTUs B_1_ H, Table 2, Figure 1). All OTUs were associated with a Family with the closest BOLD match, however OTUs C and E were similar to multiple families, as included in Table 2. OTUs D_1_ and H were the only OTUs that were identified by BOLD matches to species: *Uroobovella orri* Hirschmann (Mesostigmata: Urodinychidae) and *Diapterobates humeralis* Hermann (Sarcoptiformes: Ceratozetidae), respectively.

**Figure 1.**
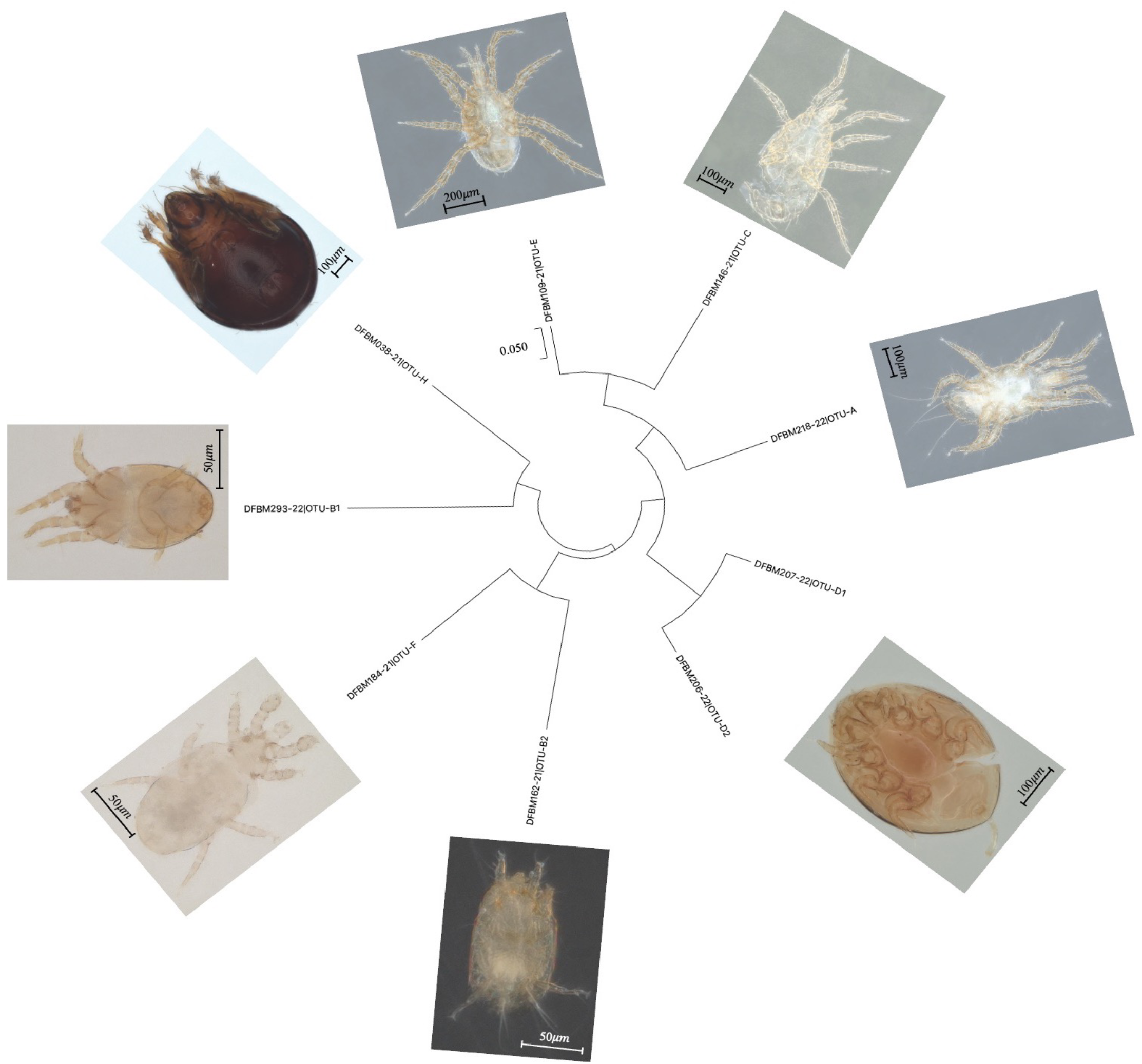
Maximum likelihood tree showing degree of relatedness between the OTUs and pictures of representatives of each OTU. Each mite branch label is designated by the OTU letter for that specimen given in Table 2.

**Table 2.** Potential taxonomic matches from DNA barcoding and prevalence of mite operational taxonomic units (OTUs) A-H from the 2021 and 2022 collections. Prevalence is the percent of Douglas-fir beetle hosts that had at least one mite of a specific OTU on their body. Abundance is the mean number of mites per beetle for an OTU. Due to similar morphology, B_1_ and B_2_ were combined in 2021 analysis, but were separated in 2022 when it became clear they were separate OTUs.

| OTU | Percent match | Order | Family | Species | Prevalence (%) |  | Abundance |  |
| --- | --- | --- | --- | --- | --- | --- | --- | --- |
|  |  |  |  |  | 2021 | 2022 | 2021 | 2022 |
| A | 92.32* | Mesostigmata | Digamasellidae | *Multidendrola elaps* sp. | 43.38 | 17.14 | 2.99 ± 0.53 | 0.33 ± 0.08 |
| B<sub>1</sub> | 98.34 | Sarcoptiformes | Hemisarcoptidae | Unknown | 96.32 | 97.86 | 69.49 ± 6.49 | 49.04 ± 4.67 |
| B<sub>2</sub> | 86.20 | Trombidiformes | Tarsonemidae | Unknown |  | 23.57 |  | 0.78 ± 0.17 |
| C | 86.21**<br>86.05 | Mesostigmata | Laelapidae,<br>Melicharidae | Unknown | 2.94 | 5.71 | 0.04 ± 0.02 | 0.09 ± 0.03 |
| D<sub>1</sub> | 99.20 | Mesostigmata | Urodinychidae | *Uroobovella orri* | 12.50 | 1.43 | 0.32 ± 0.10 | 0.01 ± 0.01 |
| D<sub>2</sub> | 95.36 | Mesostigmata | Urodinychidae | Unknown |  |  |  |  |
| E | 83.12**<br>83.12<br>82.92<br>82.59<br>82.37 | Mesostigmata | Rhodacaridae,<br>Zerconidae,<br>Laelapidae,<br>Ologamasidae,<br>Melicharidae | Unknown | 1.47 | 0.00 | 0.05 ± 0.04 | 0.00 |
| F | 84.88 | Trombidiformes | Pygmephoridae | Unknown | 6.62 | 0.00 | 0.95 ± 0.43 | 0.00 |
| G | N/A | Unknown | Unknown | Unknown | 1.47 | 6.43 | 0.03 ± 0.02 | 0.08 ± 0.03 |
| H | 98.92 | Sarcoptiformes | Ceratozetidae | *Diapterobates humeralis* | 0.74 | 0.0 | 0.01 ± 0.01 | 0.00 |
| I | N/A | Unknown | Unknown | Unknown | 0.00 | 2.14 | 0.00 | 0.02 ± 0.01 |
| J | N/A | Unknown | Unknown | Unknown | 0.00 | 0.71 | 0.00 | 0.01 ± 0.01 |

\*Initial Barcode of Life Database (BOLD) percent match of OTU A however, subsequent morphological analysis by Dr. Esmaeil Babaeian determined the identity as *Multidendrolaelaps* spp.
\*\*The National Center for Biotechnological Information BLAST program was used to determine these percent matches the associated database provided additional matches compared to the BOLD database.

OTU B_1_ was by far the most prevalent and abundant of all OTUs (Table 2). OTU B_1_ mites were consistently found grouped, often in very high numbers, in an anterior pocket on the sub-elytral surface (Figure 2). In 2022, there was a mean of 50.5 ± 4.7 mites per beetle, however when OTU B1 was removed from the analysis, the mean was only 1.3 ± 0.2 mites per beetle. Most beetles carried between 1-50 (54.9%) mites and a smaller portion carried 50-150 (33.7%) mites. Beetles were less likely to carry 150-300 mites (10.2%) and a very small portion of beetles carried 300-500 mites (1.2%). One beetle carried 456 mites, which was the maximum number that we observed. Alternatively, without OTU B_1_ in the analysis, the majority of beetles carried 0-2 (84.8%) mites and a smaller portion carried 2-14 (15.2%) mites.

**Figure 2.**
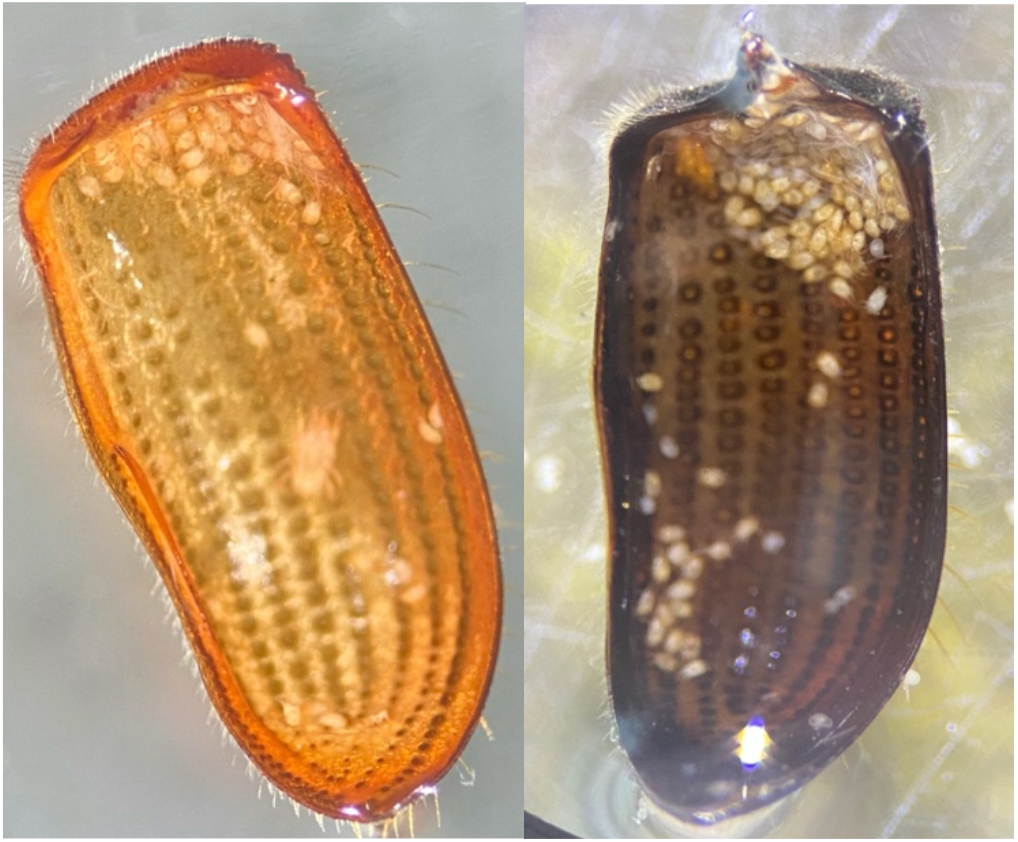
OTU B_1_ (mites clustered at the top of the images, at the anterior end of elytra) and a single representative of OTU A (in the middle of the elytron in the left image) found on the subelytral surface from two representative Douglas-fir beetles collected in this study.

Mite abundance varied throughout our sampling period (Figure 3). OTUs A and B2 were negligibly positively correlated with beetle numbers while OTU B1 was moderately, negatively correlated and C was weakly, positively correlated with the number of beetles caught (A Spearman *R* = 0.141, p-value = 0.630; B_1_ Spearman *R* = -0.410, p-value = 0.151; B_2_ Spearman *R* = 0.090, p-value = 0.770; C Spearman *R =* 0.253, p-value = 0.382). However, the high p-values associated with each spearman coefficient indicates the relationships could be due to chance. There were no differences between the mite loads of most OTUs carried by male and female beetles (B_2_ *n =* 36 females and 95 males; A, C, D_1_, D_2_, E, F, H *n =* 70 females and 191 males; Mann-Whitney p-value = 0.102 - 0.678; Table 3). OTU B_1_ was significantly differentially abundant between male and female beetles, with female beetles having significantly more associated OTU B_1_ (*n =* 36 females and 95 males; Mann-Whitney p-value = 0.0336; Table 3).

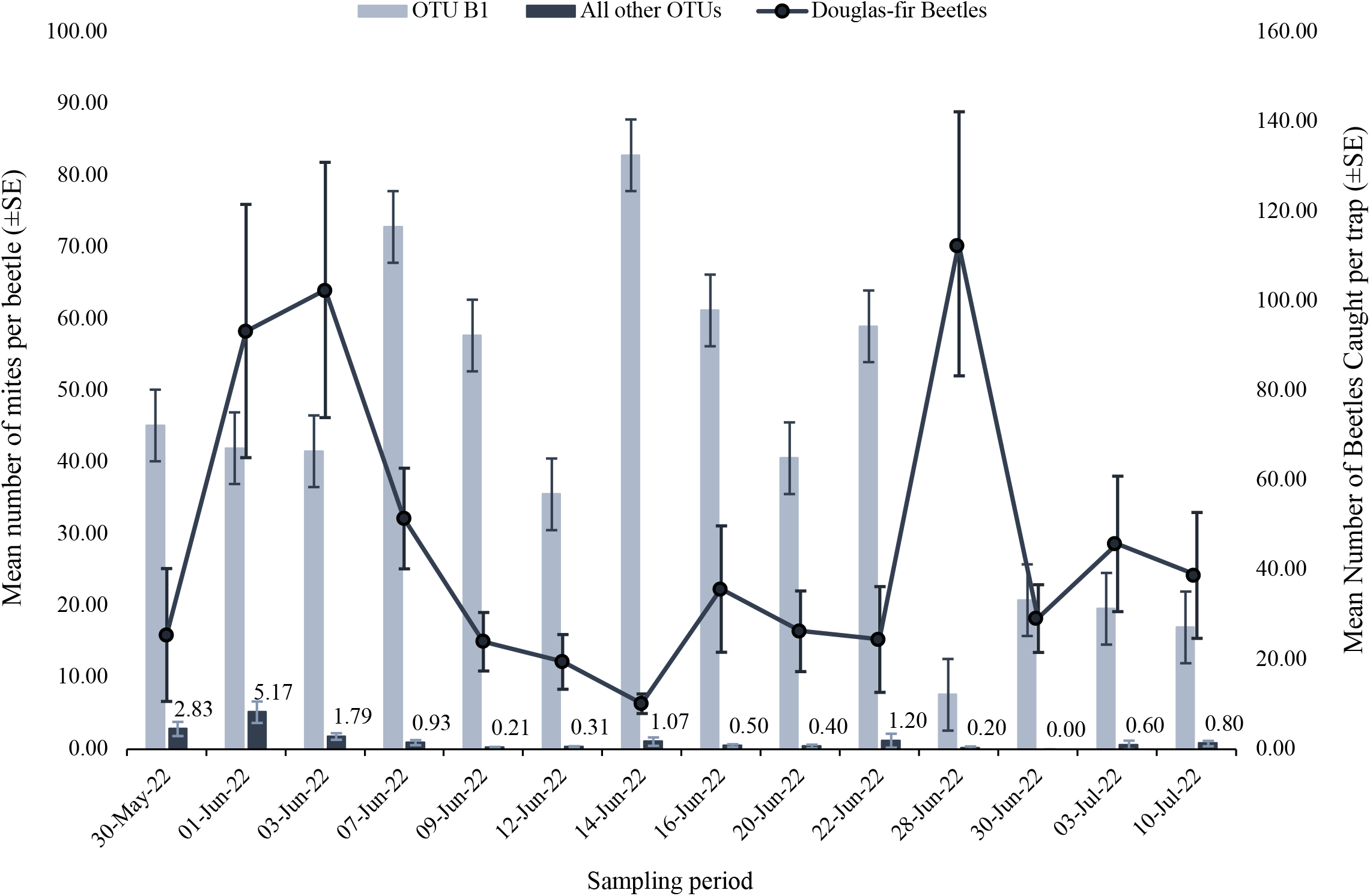

**Table 3.** Abundance (mean number of mites per beetle) of each OTU per male and female Douglas-fir beetles. OTU B_1_ (bolded) was found in significantly different abundances between beetle sexes and was found in higher numbers on female beetles.

| OTU | Abundance (mean mites/beetle ± 1 S.E.) | |
| --- | --- | --- |
|  | Male | Female |
| A | 1.24±0.35 | 1.78±0.37 |
| **B<sub>1</sub>** | **58.75±8.41** | **45.41±5.86** |
| B<sub>2</sub> | 0.69±0.22 | 0.85±0.26 |
| C | 0.06±0.03 | 0.05±0.02 |
| D<sub>1</sub> | 0.13±0.07 | 0.15±0.05 |
| D<sub>2</sub> |  |  |
| E | 0.00 | 0.04±0.03 |
| F | 0.01±0.01 | 0.67±0.30 |
| H | 0.01±0.01 | 0.00 |

## Discussion

At least nine distinct mite OTUs were associated with Douglas-fir beetle hosts as determined by successful DNA barcoding. With the additional three morphospecies that were not successfully barcoded, there are up to a dozen phoretic mite species associated with the Douglas-fir beetles in our study. Of the barcoded specimens, most were found to have BOLD matches to Families that contain species associated with other bark beetle species. Family Digamasellidae (Mesostigmata) contains *Multidendrolaelaps* spp. Hirschmann that is morphologically closely associated with *Dendrolaelaps* spp. Halbert, and for which there have been ongoing taxonomic discussions (e.g. Lindquist 1975; Huhta and Karg 2010). *Dedrolaelaps* spp. are known phoronts of several bark beetles (Coleoptera: Curculionidae) including *D. frontalis, D. ponderosae, I. typographus* and *Ips confusus* LeConte and is a predator of multiple life stages of their phoretic hosts (Hofstetter et al. 2023; Khaustov et al. 2018; Moser 1975; Vissa et al. 2020). Genera in the Family Hemisarcoptidae (Sarcoptiformes), such as *Nanacarus* spp. Oudemans, associate with tree bark, woody bracket fungi, and the galleries created by wood-boring insects – and have been found to form phoretic associations with *I. typographus* and *D. frontalis* (Hofstetter et al. 2014; Khaustov et al. 2018; Penttinen et al. 2013). The Family Tarsonemidae is a fungivorous group that contains members with fungal spore-carrying sporothecae (Moser 1985). Numerous different *Tarsonemus* spp. mites are phoretic on *I. typographus, D. ponderosae* and *D.frontalis* (Hofstetter et al. 2014; Khaustov et al. 2018; Mercado et al. 2014; Mori et al. 2011; Vissa et al. 2020). *Proctolaelaps* spp. A.Berlese (Mesostigmata: Melicharidae) and *Androlaelaps casalis* A.Berlese (Mesostigmata: Laelapidae) mites are also common associates of *I. typographus, I. confusus, D. ponderosae* and *D. frontalis* (Chaires-Grijalva et al. 2019; Hofstetter et al. 2023; Khaustov et al. 2018; Mori et al. 2011; Moser 1975; Vissa et al. 2020). *Proctolaelaps* spp. and *A. casalis* have been observed to predate on the eggs, larvae, and pupae of their associated bark beetles (Moser 1975). Mites in Family Urodinychidae develop a life stage dedicated to phoresy where they secrete an anal pedicel to attach to their hosts and they are commonly found attached to bark beetles (Knee 2012a) – the putative Urodinychidae in this study (OTUs D_1_ and D_2_) also exhibited an anal pedicel. The species match of OTU D_1_, *Uroobovella orri,* has been reported as a phoretic host generalist and is associated with multiple species of bark beetles, including the Douglas-fir beetle (Knee et al. 2012a). Pygmephoridae is a fungivorous family and certain females (phoretomorphs) will develop structures to facilitate phoretic dispersal like enlarged frontal claws (Moser and Cross 1975; Walter and Procter 2013). *Elattoma* spp. Mahunka (Trombidiformes: Pygmephoridae) have been found associated with *Ips calligraphus* (Germar) (Coleoptera: Curculionidae) and *I. typographus* (Chaires-Grijalva et al. 2019; Khaustov et al. 2018). The species match of OUT H, *Diapterobates humeralis,* has also been found in pheromone traps of bark beetles but whether they are phoretic is unconfirmed. If they are, it is likely over short distances and in response to shifting abiotic conditions (Cordes et al. 2022; Penttinen et al. 2013).

Compared to previous studies on other *Dendroctonus* spp., the mean number of mites found per Douglas-fir beetle in this study (∼50.5) was quite high. For example, the mean number of mites per *D. frontalis* beetle was found to be 4.10 and 3.96 on females and males, respectively (Moser 1976). A study on *D. ponderosae* found mean mite numbers between 0.88 to 5.50 mites per beetle while another on recently emerged *I. confusus* found an average of 18 mites per beetle (Hofstetter et al. 2023; Vissa et al. 2020). Without the hyper-abundant OTU B_1_, which was found exclusively under the anterior portion of the elytra, the mean number of mites in this study falls much closer to that recorded in other studies (∼1.3).

Studies conducted on *Hemisarcoptes* Lignières (Sarcoptiformes: Hemisarcoptidae), a relative of OTU B_1_, may shed light onto the relationship between Douglas-fir beetles and the hyper-abundant OTU B_1_. All non-phoretic lifestages of *Hemisarcoptes* are predators of scale insects (Hemiptera: Diaspididae), but in their phoretic stage, referred to as a hypopus, the mites lose functional mouthparts and develop ventral suckers to attach to the sub-elytral surface of their *Chilocorus* spp. Leach (Coleoptera: Coccinellidae) hosts. One study reported the distribution of *Hemisarcoptes* under their host elytra and noted that they were non-randomly attached on the epipleural margin of the subelytral surface and in very high numbers (400-800 per beetle), like the non-random distribution of OTU B_1_ also in high numbers (Houck 1999). The study related the attachment patterns of *Hemisarcoptes* to the distribution of spines along the host’s subelytral surface, which the author hypothesized could damage the ventral surface of *Hemisarcoptes* (Houck 1999).

The distribution of subelytral spines seem to dictate where mites are able to attach underneath their host elytra. Further studies have found that not only can *Hemisarcoptes* acquire beetle hemolymph through a vestigial anal opening, but they are also capable of passing material back to the beetle, meaning there is a bi-directional flow of materials at the attachment site of *Hemisarcoptes* (Holte et al. 2001; Houck 1994; Houck and Cohen 1995). No further studies have been conducted into the purpose of sharing materials between phoronts and hosts, though speculations include potentially communicating through sharing hormones or enzymes as well as a method of receiving amino acids or sugars (Houck 1999). Due to not having an oral opening, *Hemisarcoptes* may also excrete harmful metabolic waste into their beetle hosts (Holte et al, 2001). In either case, having such high loads of mites that are able to remove and input materials could impact the success of hosts, but it is not clear yet if the fitness consequences on the host are positive, negative, or neutral.

If Douglas-fir beetles also have some control over where these mites can attach, through the distribution of subelytral structures or other anatomical-behavioral influences, they may have developed a pattern of distribution that minimizes any hindrances to mobility and other impacts on success. Elytral adaptations associated with symbionts are not new in Douglas-fir beetles, which contain elytral pits that carry beneficial fungal spores similar to the mycangia of other bark beetles (Lewinsohn et al. 1994). Other bark beetles have pronotal mycangia that contain glands that interact with the fungi they carry (Pechanova et al. 2008). It is possible there is some other co-evolved relationship between the Douglas-fir beetle and OTU B_1_ besides phoresy, or perhaps phoresy is only one aspect of the interaction. However, further research is required into the presence of the sub-elytral anatomy and semiochemical environment in Douglas-fir beetles and the possibility of an analogous bi-directional flow of material between the organisms for physiological or communicative purposes.

Throughout the collection period, Douglas-fir beetles emerged in highest numbers in late May and early June and subsequently declined until spiking in numbers on a single date near the end of collection (Figure 3). OTU B_1_ was most numerous on Douglas-fir beetles during much of the collection period before sharply declining on the late emerging beetles (Figure 3). Other OTUs were most abundant on the early flying beetles before mostly declining throughout the rest of the sampling period (Figure 3). Previous studies have noted that mite abundances tended to peak at similar times as the flight peaks of their bark beetle hosts and during the highest abundances of their coleopteran host (Knee et al. 2012b; Paraschiv et al. 2018). This is mostly supported by OTUs A, B_2_ and C being negligibly and moderately positively correlated with the number of beetles. Hofstetter et al. (2023) found that species within families Digamasellidae and Tarsonemidae (related to OTUs A and B_2_) were most abundant on early emerging beetles. Interestingly, OTU B_1_ was moderately negatively correlated with the beetle numbers, indicating it may have an adverse impact on Douglas-fir beetle populations. However, due to the high p-values associated with the Spearman coefficients, we were unable to determine if the resulting relationships between the number of beetles caught and the number of mites per beetle are due to chance.

All except one OTU were found equally on male and female beetles meaning there is no sex bias for most OTUs. Knee et al. (2012b) hypothesized that on cerambycid beetles, due to a female beetle’s proximity to the egg niche where males do not enter, mites may have an advantage entering the beetle’s gallery when associated with a female. However, in the case of the Douglas-fir beetle, both males and females construct the egg gallery and therefore, mites attached to both sexes would have equal opportunity to enter the tree. This finding is supported by other studies of the potential sex bias of phoretic mites on other bark beetles, which showed no sex bias (Mori et al. 2011; Moser 1976; Paraschiv et al. 2018). That being said, OTU B_1_ was found in significantly higher numbers on female Douglas-fir beetles. Houck (1999) noted that larger body sizes in coccinellid hosts provided more space for mite attachment. Douglas-fir beetle females are generally larger than males and so higher numbers of the most prevalent mite may simply be a matter of more attachment space.

This study is one of the first to document the diversity, phenology, host preference and attachment location of the phoretic mite assemblage of the Douglas-fir beetle. The mite assemblage includes at least nine distinct OTUs coming from multiple different families with unique life histories and potential impacts on the Douglas-fir beetle. Particularly, we hypothesize the potential importance of the hyper-abundant OTU B_1_ to Douglas-fir beetles during their adult host colonization phase, and we suggest that this phenomenon deserves further research in this insect and in analogous situations with other Scolytinae.

## Acknowledgements

We would like to thank Mackenzie Howse, Claire Paillard, Anne Robinson, Margaret Browning, Steve Browning, and Chantal Presse for their help in the field. We also thank Dr. Aija White and Dr. Heather Bryan for their discussions throughout this study. This project was supported by a Natural Sciences and Engineering Research Council of Canada Undergraduate Student Research Award (L-A. B.) and Discovery Grant (DPWH).

## Literature cited

1. Atkins, M. D. 1960. A Study of the Flight of the Douglas-Fir Beetle Dendroctonus pseudotsugae Hopk. (Coleoptera : Scolytidae) II. Flight Movements. The Canadian Entomologist, 92(12): 941–954. https://doi.org/10.4039/Ent92941-12

2. Bartlow, A. W., and Agosta, S. J. 2021. Phoresy in animals: review and synthesis of a common but understudied mode of dispersal. Biological Reviews, 96(1): 223–246. https://doi.org/10.1111/brv.12654

3. Bentz, B. J., Régnière, J., Fettig, C. J., Hansen, E. M., Hayes, J. L., Hicke, J. A., Kelsey, R. G., Negrón, J. F., and Seybold, S. J. 2010. Climate change and bark beetles of the western United States and Canada: Direct and indirect effects. BioScience, 60(8): 602–613. https://doi.org/10.1525/bio.2010.60.8.6

4. Chaires-Grijalva, M. P., Estrada-Venegas, E. G., Quiroz-Ibáñez, I. F., Equihua-Martínez, A., Moser, J. C., and Blomquist, S. R. 2019. Acarine biodiversity associated with bark beetles in Mexico. Acarological Studies, 1(2): 152–160.

5. Cole, H., Andrus, R. A., Butkiewicz, C., Rodman, K. C., Santiago, O., Tutland, N. J., Waupochick, A., and Hart, S. 2022. Outbreaks of Douglas-Fir Beetle Follow Western Spruce Budworm Defoliation in the Southern Rocky Mountains, USA. Forests, 13(3): 371. https://doi.org/10.3390/f13030371

6. Cordes, P., Maraun, M., and Schaefer, I. 2022. Dispersal patterns of oribatid mites across habitats and seasons. Experimental and Applied Acarology, 86: 173–187. https://doi.org/10.1007/s10493-022-00686-y

7. Edgar, R. C. 2004. MUSCLE: a multiple sequence alignment method with reduced time and space complexity. BMC Bioinformatics, 5:1–19. https://doi.org/10.1186/1471-2105-5-113

8. Hofstetter, R. W., Klepzig, K. D., Moser, J. C., and Ayres, M. P. 2006. Seasonal Dynamics of Mites and Fungi and Their Interaction with Southern Pine Beetle. Environmental Entomology, 35(1): 22–30. https://doi.org/10.1603/0046-225x-35.1.22

9. Hofstetter, R.W., Moser, J.C., Blomquist, S. 2014. Mites associated with bark beetles and their hyperphoretic ophiostomatoid fungi. CBS Biodiversity Series, 12: 165–176.

10. Hofstetter, R. W., Dinkins-Bookwalter, J., Davis, T. S., and Klepzig, K. D. 2015. Symbiotic Associations of Bark Beetles. *In* Bark Beetles: Biology and Ecology of Native and Invasive Species. *Edited by* F. E. Vega and R. W. Hofstetter. Academic Press. Pp. 209–245. https://doi.org/10.1016/B978-0-12-417156-5.00006-X.

11. Hofstetter, E. M., Knee, W. H., and Khaustov, A. A. 2023. Phoretic mite assemblage of the pinyon pine beetle, Ips confusus (Curculionidae: Scolytinae), in Arizona. Acarologia, 63(2): 480–490. https://doi.org/10.24349/upy5-taez

12. Holte, A. E., Houck, M. A., and Collie, N. L. 2001. Potential role of parasitism in the evolution of mutualism in astigmatid mites: Hemisarcoptes cooremani as a model. Experimental and Applied Acarology, 25(2): 97–107. https://doi.org/10.1023/a:1010655610575

13. Houck, M. A. 1994. Adaptation and Transition into Parasitism from Commensalism: A Phoretic Model. In Mites. Edited by M.A. Houck. Springer, Boston, Massachusetts, United States. Pp. 252–281. https://doi.org/10.1007/978-1-4615-2389-5_10

14. Houck, M. A. 1999. Phoresy by Hemisarcoptes (Acari: Hemisarcoptidae) on Chilocorus (Coleoptera: Coccinellidae): influence of subelytral ultrastructure. *In* Ecology and Evolution of the Acari. *Edited by* J. Bruin, L.P.S van der Geest, M.W. Sabelis. Springer, Dordrecht, The Netherlands. Pp. 303–321. https://doi.org/10.1007/978-94-017-1343-6_21

15. Houck, M. A., and Cohen, A. C. 1995. The potential role of phoresy in the evolution of parasitism: radiolabelling (tritium) evidence from an astigmatid mite. International Journal of Acarology, 19(12): 677–694. https://doi.org/10.1007/bf00052079

16. Huhta, V., and Karg, W. 2010. Ten new species in genera Hypoaspis (s. lat.) Canestrini, 1884, Dendrolaelaps (s. lat.) Halbert, 1915, and Ameroseius Berlese, 1903 (Acari: Gamasina) from Finland. Soil Organisms, 82(3): 325–349.

17. Kelley, S. T., and Farrell, B. D. (1998). Is Specialization a Dead End? The Phylogeny of Host Use in Dendroctonus Bark Beetles (Scolytidae). Evolution, 52(6), 1731–1743. https://doi.org/10.2307/2411346

18. Khaustov, A. A., Klimov, P. B., Trach, V. A., Bobylev, A. N., Salavatulin, V. M., Khaustov, V. A., and Tolstikov, A. V. 2018. Review of Mites (Acari) Associated with the European Spruce Bark Beetle, Ips Typographus (Coleoptera: Curculionidae: Scolytinae) in Asian Russia. Acarina, 26(1): 3–79. https://doi.org/10.21684/0132-8077-2018-26-1-3-79

19. Knee, W., Beaulieu, F., Skevington, J. H., Kelso, S., Cognato, A. I., and Forbes, M. R. 2012a. Species Boundaries and Host Range of Tortoise Mites (Uropodoidea) Phoretic on Bark Beetles (Scolytinae), Using Morphometric and Molecular Markers. PLoS ONE, 7(10): e47243. https://doi.org/10.1371/journal.pone.0047243

20. Knee, W., Hartzenberg, T., Forbes, M. R., and Beaulieu, F. 2012b. The natural history of mites (Acari: Mesostigmata) associated with the white-spotted sawyer beetle (Coleoptera: Cerambycidae): diversity, phenology, host attachment, and sex bias. The Canadian Entomologist, 144(5): 711–719. https://doi.org/10.4039/tce.2012.57

21. Lewinsohn, D., Lewinsohn, E., Bertagnolli, C. L., and Patridge, A. D. 1994. Blue-stain fungi and their transport structures on the Douglas-fir beetle. Canadian Journal of Forest Research, 24(11): 2275–2283. https://doi.org/10.1139/x94-292

22. Lindgren, B.S. 1983. A multiple funnel trap for scolytid beetles (Coleoptera). The Canadian Entomologist. 115(3): 299–302.

23. Lindquist, E. E. 1975. *Digamasellus* Berlese, 1905, and *Dendrolaelaps* Halbert, 1915, with Descriptions of New Taxa of Digamasellidae (Acarina: Mesostigmata). Canadian Entomologist, 107(1): 1–43. https://doi.org/10.4039/ent1071-1

24. Lombardero, M. J., Ayres, M. P., Hofstetter, R. W., Moser, J. C., and Lepzig, K. D. 2003. Strong indirect interactions of Tarsonemus mites (Acarina: Tarsonemidae) and Dendroctonus frontalis (Coleoptera: Scolytidae). Oikos, 102(2): 243–252. https://doi.org/10.1034/j.1600-0706.2003.12599.x

25. Lyon, R. L. 1958. A Useful Secondary Sex Character in Dendroctonus Bark Beetles. The Canadian Entomologist, 90(10): 582–584. https://doi.org/10.4039/ent90582-10

26. Mercado, J. E., Hofstetter, R. W., Reboletti, D. M., and Negrón, J. F. 2014. Phoretic Symbionts of the Mountain Pine Beetle (Dendroctonus ponderosae Hopkins). Forest Science, 60(3): 512–526. https://doi.org/10.5849/forsci.13-045

27. Ministry of Forests, Lands, Natural Resource Operations and Rural Development. 2021. Aerial Overview Survey Summary Reports - Province of British Columbia. Government of British Columbia. Available from https://www2.gov.bc.ca/gov/content/industry/forestry/managing-our-forest-resources/forest-health/aerial-overview-surveys/summary-reports [accessed 01 June 2023].

28. Mori, B. A., Proctor, H. C., Walter, D. E., and Evenden, M. L. 2011. Phoretic mite associates of mountain pine beetle at the leading edge of an infestation in northwestern Alberta, Canada. The Canadian Entomologist, 143(1): 44–55. https://doi.org/10.4039/n10-043

29. Moser, J. C. 1975. Mite Predators of the Southern Pine Beetle. Annals of the Entomological Society of America, 68(6): 1113–1116. https://doi.org/10.1093/aesa/68.6.1113

30. Moser, J. C. 1976. Surveying Mites (Acarina) Phoretic on the Southern Pine Beetle (Coleoptera: Scolytidae) with Sticky Traps. The Canadian Entomologist, 108(8): 809–813. https://doi.org/10.4039/ent108809-8

31. Moser, J. C. 1985. Use of sporothecae by phoretic Tarsonemus mites to transport ascospores of coniferous bluestain fungi. Transactions of the British Mycological Society, 84(4): 750– 753. https://doi.org/10.1016/s0007-1536(85)80138-8

32. Moser, J. C., and Cross, E. A. 1975. Phoretomorph: A New Phoretic Phase Unique to the Pyemotidae (Acarina: Tarsonemoidea). Annals of the Entomological Society of America, 68(5): 820–822. https://doi.org/10.1093/aesa/68.5.820

33. Paraschiv, M., Martínez-Ruiz, C., and Fernández, M. M. 2018. Dynamic associations between Ips sexdentatus (Coleoptera: Scolytinae) and its phoretic mites in a Pinus pinaster forest in northwest Spain. Experimental and Applied Acarology, 75(3): 369–381. https://doi.org/10.1007/s10493-018-0272-9

34. Pechanova, O., Stone, W., Monroe, W. S., Nebeker, T. E., Klepzig, K. D., and Yuceer, C. 2008. Global and comparative protein profiles of the pronotum of the southern pine beetle, Dendroctonus frontalis. Insect Molecular Biology, 17(3): 261–277. https://doi.org/10.1111/j.1365-2583.2008.00801.x

35. Penttinen, R., Viiri, H., and Moser, J. 2013. The mites (Acari) associated with bark beetles in the Koli National Park in Finland. Acarologia, 53(1): 3–15. https://doi.org/10.1051/acarologia/20132074

36. Raffa, K. F., Aukema, B. H., Bentz, B. J., Carroll, A. L., Hicke, J. A., Turner, M. G., and Romme, W. H. 2008. Cross-scale Drivers of Natural Disturbances Prone to Anthropogenic Amplification: The Dynamics of Bark Beetle Eruptions. BioScience, 58(6): 501–517. https://doi.org/10.1641/b580607

37. Ratnasingham, S., and Hebert, P. D. N. 2007. BARCODING: bold: The Barcode of Life Data System (http://www.barcodinglife.org). Molecular Ecology Notes, 7(3): 355–364. https://doi.org/10.1111/j.1471-8286.2007.01678.x

38. Rocha, S., Pozo-Velázquez, E., Faroni, L., and Guedes, R. 2009. Phoretic load of the parasitic mite Acarophenax lacunatus (Cross and Krantz) (Prostigmata: Acarophenacidae) affecting mobility and flight take-off of Rhyzopertha dominica (F.) (Coleoptera: Bostrichidae). Journal of Stored Products Research, 45(4): 267–271. https://doi.org/10.1016/j.jspr.2009.05.001

39. Ruiz, E. A., Rinehart, J. E., Hayes, J. L., and Zuñiga, G. 2010. Historical Demography and Phylogeography of a Specialist Bark Beetle, Dendroctonus pseudotsugae Hopkins (Curculionidae: Scolytinae). Environmental Entomology, 39(5): 1685–1697. https://doi.org/10.1603/en09339

40. Stark, R. W. 1982. Generalized Ecology and Life Cycle of Bark Beetles. In Bark Beetles in North American Conifers: A System for the Study of Evolutionary Biology. Edited by J. B. Mitton and K. B. Sturgeon. University of Texas Press. Pp. 21–45

41. Stecher, G., Tamura, K., and Kumar, S. 2020. Molecular Evolutionary Genetics Analysis (MEGA) for macOS. Molecular Biology and Evolution, 37(4): 1237–1239. https://doi.org/10.1093/molbev/msz312

42. Vissa, S., Mercado, J. E., Malesky, D., Uhey, D. A., Mori, B. A., Knee, W., Evenden, M. L., and Hofstetter, R. W. 2020. Patterns of Diversity in the Symbiotic Mite Assemblage of the Mountain Pine Beetle, Dendroctonus Ponderosae Hopkins. Forests, 11(10): 1102. https://doi.org/10.3390/f11101102

43. Walter, D. E., and Proctor, H. C. 2013. Mites: Ecology, Evolution and Behaviour: Life at a Microscale. Springer, Dordrecht, The Netherlands.

